# MRGM: a mouse reference gut microbiome reveals a large functional discrepancy for gut bacteria of the same genus between mice and humans

**DOI:** 10.1101/2021.10.24.465599

**Authors:** Nayeon Kim, Chan Yeong Kim, Sunmo Yang, Dongjin Park, Sang-Jun Ha, Insuk Lee

## Abstract

The gut microbiome is associated with human diseases and interacts with dietary components and drugs. *In vivo* mouse models may be effective for studying diet and drug effects on the gut microbiome. We constructed a mouse reference gut microbiome (MRGM, https://www.mbiomenet.org/MRGM/) that includes newly-assembled genomes from 878 metagenomes. Leveraging samples with ultra-deep metagenomic sequencing (>130 million read pairs), we demonstrated quality improvement in assembled genomes for mouse gut microbes as sequencing depth increased. MRGM provides a catalog of 46,267 non-redundant genomes with ≥70% completeness and ≤5% contamination comprising 1,689 representative bacterial species and 15.2 million non-redundant proteins. Importantly, MRGM significantly improved the taxonomic classification rate of sequencing reads from mouse fecal samples compared to previous databases. Using MRGM, we determined that reliable low-abundance taxa profiles of the mouse gut microbiome require sequencing >10 million reads. Despite the high overall functional similarity of the mouse and human gut microbiomes, only ~10% of MRGM species are shared with the human gut microbiome. Although ~80% of MRGM genera are present in the human gut microbiome, ~70% of the shared genera have <40% of core gene content for the respective genus with human counterparts. These suggest that although metabolic processes of the human gut microbiome largely occur in the mouse gut microbiome, functional translations between them according to genus-level taxonomic commonality require caution.

**Key Points:**

1. MRGM provides 46,267 genomes comprising 1,689 bacterial species of mouse gut microbiome.
2. Despite high overlap of genera, functional discrepancy between mouse and human gut microbiota is large.
3. Lineage-specific markers underestimate the completeness of assembled genomes for uncharacterized taxa.

## INTRODUCTION

During the past several decades, numerous studies have provided accumulated evidence of an association between the gut microbiome and human diseases (1). These findings inspired the idea of microbiome medicine, where the gut microbiome can be leveraged as a biomarker and therapeutic target to improve disease conditions (2–4). As the human reference genome has opened the era of genome medicine, the human reference gut microbiome will allow microbiome medicine to become feasible (5–7). Recently, a number of reference gut microbiomes specific for humans have been published (8–10), and the latest of these microbiomes provides ~232k non-redundant genomes comprising 5,414 prokaryotic species (10). These new catalogs exhibited substantially improved taxonomic and functional classification of metagenomic sequence reads compared to those of the previously used standard reference catalog. Mice can provide *in vivo* models for human gut microbiota research (11), and based on this, establishing comprehensive catalogs of mouse gut microbial genomes will also facilitate our understanding of the human gut microbiome.

To date, only two large-scale culture-based studies, miBC and mGMB, have reported 76 and 126 isolated species, respectively, from the mouse gut environment (12,13). Furthermore, the collection of metagenome-assembled genomes (MAGs) for the mouse gut microbiome lags behind that of the human gut microbiome. *De novo* genome assembly from gut metagenome sequencing samples obtained from 184 mice cataloged ~2.6 million non-redundant protein coding genes for 541 microbial species (14). A more recently published reference gut microbiome for mice, iMGMC, cataloged 18,203 MAGs possessing medium quality (completeness ≥ 50% and contamination ≤ 10%), including those from co-assembly (all-in-one) (15). The co-assembly approach allows for the retrieval of low-abundance genomes by complementing sequence reads from different samples. However, this method is susceptible to mismatching of genomic fragments from different strains of the prevalent species (16). Therefore, strain-level analysis requires MAGs derived only from a single-sample assembly.

Here, we present the mouse reference gut microbiome (MRGM) that provides 46,267 non-redundant genomes with a completeness of ≥ 70% and a contamination of ≤ 5% and is comprised of 1,689 representative prokaryotic species and ~15.2 million encoded proteins. In addition to the consolidated isolated genomes and MAGs available from the public repository, we deposited novel MAGs by conducting *de novo* assembly using public whole metagenomic sequencing (WMS) data from 838 mouse fecal or cecal samples and newly-generated WMS data from 40 mouse fecal samples. To improve the usability of the database, we provide a web server (https://www.mbiomenet.org/MRGM/) where users can browse the cataloged genomes with all relevant information.

Taxonomic and functional discrepancies in the gut microbiome between mice and humans will likely cause pitfalls in human gut microbiome research that is based on mouse models (11). Using MRGM, we performed taxonomic and functional comparisons of the gut microbiome between mice and humans. The information obtained from this analysis will provide useful guidelines for translating metagenomic research from mouse models into the human gut microbiome. Consistent with previous reports (14), the overall functional capacity of the human gut microbiome is largely present in the mouse gut microbiome; however, only ~10% of bacterial species from the mouse gut are shared in humans. Information from mouse gut microbiome research is often transferred to the human gut microbiome via shared genera between these microbiomes. Therefore, we evaluated the taxonomic and functional overlap between mouse and human gut microbiomes at the genus level. We observed that although ~80% of mouse gut bacterial genera are shared in the human gut microbiome, ~70% of the shared genera have <40% overlap of core gene content for the respective genus. These results suggest difficulties regarding the study of the human gut microbiome when using the taxonomic profiles of the mouse gut microbiome.

## MATERIALS AND METHODS

### Whole metagenome sequencing of mouse fecal samples

We generated WMS data for 40 fecal samples obtained from 40 C57BL/6J mice with a sequencing depth of 12-47 Gbp. Library preparation was performed according to the TruSeq Nano DNA Library Preparation Guide. First, 100 ng of DNA was cleaved using a Covaris LE200 Focused-ultrasonicator. After conversion into blunt-end DNA, DNA fragments of approximately 350 bp were selected. DNA fragments were ligated using an indexing adapter. DNA fragments were amplified by eight cycles of PCR. We estimated the quality and band size of libraries using D1000 Screen Tapes (Agilent) on a Tapestation 4200 (Agilent). The libraries were quantified using KAPA Library Quantification kits for Illumina Sequencing platforms according to the qPCR Quantification Protocol Guide (KAPA BIOSYSTEMS, #KK4854). Finally, the libraries were sequenced using the HiSeq 4000 platform (Illumina, San Diego, USA).

### Metagenome *de novo* assembly and binning

In addition to the in-house WMS data generated from 40 mouse fecal samples, we collected public WMS data for 838 samples that were recently deposited but not included in the previous catalogs. Samples from mice that received fecal microbiota transplantation from humans were excluded. We removed adapters and low-quality bases from the sequence reads using Trimmomatic v0.39 (17). Next, we removed host-contaminated sequences using Bowtie2 v2.3.5 (18) by aligning sequence reads to the mouse genome GRCm39. High-quality reads were assembled into contigs using MetaSPAdes v3.14.1 (19) or MEGAHIT v1.2.9 (20). MetaSPAdes was used as the default assembly software. If the assembly could not be completed by MetaSPAdes for unknown reasons or if WMS data were based on single-end sequencing, the assembly was performed using MEGAHIT.

Binning, grouping contigs into the genome bins, was performed using an ensemble approach. Prior to binning, the sequence reads were aligned to the contigs using Bowtie2. Next, three different binning tools, specifically MetaBAT2 v2.15 (21), MaxBIN2 v2.2.7 (22), and CONCOCT v1.1.0 (23), were used to generate genome bins separately. We set the minimum contig length to 1,000 for MaxBin2 and CONCOCT and to 1,500 for MetaBAT2. Next, we consolidated the binning results using the bin refinement module of MetaWRAP v1.3.2 (24) that estimated the quality of genome bins using CheckM (25) and selected the best result.

### Public MAGs and isolated genomes for mouse gut microbiome

We collected publicly available MAGs and isolated genomes for the mouse gut microbiome from MMGC (26) (as of February 2021) and from iMGMC (15) (only sample-specific MAGs were taken). We also collected isolated genomes from miBC (12), mGMB (13), and PATRIC (27).

### Genome quality assessment

We assessed the completeness and contamination of genome bins using the CheckM v1.1.3 (25) lineage-specific workflow that estimates genome quality based on completeness and contamination using lineage-specific marker gene sets. Genomes were further assessed using GUNC v1.0.1 (28), a software program that detects genome chimerism and reports clade separation scores (CSS). We filtered out genomes exhibiting CSS > 0.45, as this is the default threshold of the software.

### Generation of genomic species clusters

We clustered genomes into species-level genome bins (SGBs) using a two-step iterative procedure that included fast preliminary clustering followed by more accurate secondary clustering. Preliminary clustering was performed according to average-linkage hierarchical clustering at a cutoff of 0.2 using the Mash v2.2.2 (29) distance. The mash distance was calculated for all pairwise distances between genomes using a sketch size of 10,000. Mash can calculate all pairwise distances between genomes relatively quickly. However, when the coverage of the genome is low, the accuracy will be reduced. We complemented the low accuracy of the initial clusters through refined secondary clustering using the ANImf of dRep (16). ANImf was calculated for every pair of genomes within each initial cluster. To avoid overestimation of average nucleotide identity (ANI) by local alignment, a minimum coverage threshold was applied for each pair. The coverage cutoff of genome A and genome B was determined at min (0.8, (*Completeness of genome A* * *Completeness of genome B*)). Genomes were clustered by ≥95% ANI, as this is equivalent to ANI among the same bacterial species (30). In each cluster, we chose a representative genome with the highest score of the genome intactness score according to the equation S = *Completeness* - 5 × Contamination + 0.5 × *log*_10_ *N*_50_. Next, using the selected representative genomes, we clustered genomes iteratively using a preceding two-step clustering until they were no longer clustered. When we counted the conspecific genomes for each species, and we clustered average-linkage hierarchical clustering at a cutoff of 0.001 using the Mash distance (Mash ANI 99.9%). We also selected only one genome per species per sample.

### Taxonomic annotation and phylogenetic tree construction

We conducted taxonomic annotation for species-representative genomes based on the Genome Taxonomy Database (GTDB) R95 (31). We used GTDB-Tk v1.4.1 (32) to classify query genomes to GTDB taxa based on its reference tree using 120 bacterial and 122 archaeal marker genes. GTDB-Tk aligned the marker genes and generated multiple sequence alignments of these genes for each species-representative genome. We inferred a phylogenetic tree using IQ-TREE v.2.0.3 (33) based on multiple sequence alignment of the concatenated sequences of 120 bacterial marker genes. For visualization of the phylogenetic tree, we used iTOL v5 (34).

### Identifying 16S rRNA sequences from reference gut bacterial genomes

We predicted the 16S rRNA sequences from the representative species genomes using barrnap v0.9 (35). We lowered the default threshold of the e-value from 1e-06 to 1e-05. As highly conserved rRNA genes are difficult to predict from MAGs that are generated from short-read sequences, we used all non-redundant genomes rather than only representative genomes. We cataloged the 16S rRNA sequences from representative genomes by default. If representative genomes possessed no 16S rRNA sequence, we cataloged this sequence from the longest genomes of the species cluster.

### Cataloging mouse gut microbial proteins and their functional annotation

To catalog mouse gut microbial proteins, we predicted coding sequences (CDS) from genomes using Prodigal v2.6.3 (36) with the parameter “-c -m -p single”. To avoid redundancy, we performed clustering of protein sequences for 100% similarity using linclust (37) from MMseq2 v.13.45111 (38). We performed linclust with parameter “--cov-mode 1–c 0.8 --cluster-mode 2 --kmer-per-seq 80 --min-seq-id 1”. Sequentially, we performed clustering at 95%, 90%, 70%, and 50% sequence similarity by altering the parameter --min-seq-id with 0.95, 0.90, 0.70, and 0.5. Cataloged mouse gut microbial proteins were annotated using eggNOG-mapper v.2.1.2 (39) with the parameter “-m diamond” to provide functional annotations based on KEGG pathways/modules/orthologs (40), Gene Ontology (41), CAZy terms (42), and enzyme commission (EC) numbers.

### Assessment of sequencing depth effect on MAG assembly

To test whether MAG assembly is improved by increased sequencing depth, we generated simulated datasets for various sequencing depths. We selected 10 WMS samples possessing the largest sequencing depths out of the 40 samples according to ultra-deep sequencing (>40 Gbp or >130 million read pairs). For each sample, 0.5, 2.5, 5, 10, 20, 40, 80, and 125 million read pairs (i.e., 150 Mbp, 750 Mbp, 1.5 Gbp, 3 Gbp, 6 Gbp, 12 Gbp, and 37.5 Gbp in total sequencing depth for read pair length of 300 bp) were randomly selected. Next, 80 simulated datasets (10 samplesⅹ8 depths) were assembled using the same in-house pipeline. We tested whether the quality of the conspecific genomes was improved as the sequencing depth was increased. We performed average-linkage hierarchical clustering at a cutoff of 0.1 using the Mash distance (Mash ANI 90%). Thereafter, if the MAGs from the same sample but at different depths were clustered, the MAGs were considered to be conspecific genomes. We compared the quality of the conspecific MAGs from adjacent sequencing depths.

We also tested whether deep sequencing aided in the assembly of MAGs for low-abundance taxa. All 878 WMS samples used for MRGM construction were divided into two groups that included >10 Gbp and <10 Gbp WMS sample groups. We measured the relative abundance of taxa whose MAGs were assembled specifically from each group of samples and did not exist in iMGMC or MMGC (i.e., newly assembled in MRGM). The relative abundances of taxa in 10 WMS samples obtained from PRJNA603829 (https://www.ebi.ac.uk/ena/browser/view/PRJNA603829) that were not included in the MRGM construction were estimated using Kraken2.

### Evaluation of the taxonomic classification rates of the metagenomic sequencing reads

We compared taxonomic classification rates according to different reference gut microbial genome databases using Kraken2 v2.0.8-beta (43). The standard Kraken2 database was downloaded using “kraken2-build --standard”. In addition to the standard Kraken2 database that is based on RefSeq prokaryotic genomes, we generated a custom Kraken2 database for iMGMC (15) using 1,296 representative mMAGs that were downloaded from GitHub (https://github.com/tillrobin/iMGMC). To generate custom a Kraken2 database for MRGM, we used the representative genomes for each species cluster. We downloaded a pre-built custom Kraken2 database for representative species genomes of MMGC (26) as of February 2021. For comparison, we also included a custom Kraken2 database for HRGM (10) that is available from a web server (https://www.mbiomenet.org/HRGM/). If the given representative species genome possessed no GTDB taxon, we assigned a hypothetical taxon. For example, if a genome was annotated at the family level only, we assigned an additional ID to the genus and species levels. Certain genomes were annotated to the same species despite differences in ANI ≥ 95. In this case, we separated the genomes into different species by assigning an additional ID.

For the assessment of taxonomic classification rates, we used WMS data obtained from PRJEB37572 (44) and PRJNA730805 (45) that were not used for the construction of any of the genome catalogs. PRJEB37572 and PRJNA730805 contained 86 samples of single-end sequencing and 30 samples of paired-end sequencing, respectively. Prior to taxonomic classification, we also pre-processed the WMS data using Trimmomatic and Bowtie2. For paired-end sequencing samples, we performed Kraken2 with the option “paired”. The taxonomic classification rate was calculated based on the percentage of reads that were classified.

### Assessment of the sequencing depth effect on taxonomic profiling

Taxonomic features at the domain, phylum, class, order, family, genus, and species levels were stratified into eight groups according to mean relative abundance (ranging from 1e-7 to 1 with every ten-fold increase) using 10 WMS samples possessing the largest sequencing depths (>80 Gbp or >260 million read pairs) of PRJNA603829 (https://www.ebi.ac.uk/ena/browser/view/PRJNA603829) that were not used for MRGM. For each of the 10 WMS samples, simulated datasets were generated for 10 different depth ranges that included 0.5, 1, 2.5, 5, 10, 20, 40, 60, 80, and 125 million read pairs. Taxonomic profiles for the 100 simulated datasets were generated using the custom Kraken2 database based on MRGM. As the group for mean relative abundance of <1e-7 included only three taxonomic features, we combined it with the group for mean relative abundance of <1e-6. We then calculated the Pearson correlation coefficient (*PCC*) and Spearman correlation coefficient (*SCC*) between the taxonomic profiles of relative abundance at different sequencing depths for each level of taxonomic features.

### Comparison of core genes for the same bacterial genus between humans and mice

Protein clusters for 50% similarity of MRGM and HRGM were used to compare core genes for the same bacterial genus between humans and mice. Based on the GTDB taxonomic annotation, MRGM contains 272 genera. We conducted a comparison of 220 genera that existed in both MRGM and HRGM. Genes that were conserved among ≥90% of non-redundant genomes of each genus were defined as ‘genus core’ genes. As we compared core genes for the same bacterial genus between humans and mice, we grouped core genes of humans and mice together according to 50% sequence identity using linclust with parameter ‘--min-seq-id 0.5’. For the same genus of gut bacterial genomes, we calculated the proportion of shared genus core genes between humans and mice.

## RESULTS

### Cataloging genomes and proteins for 1,689 prokaryotic species in the mouse gut

Our in-house pipeline for cataloging the microbial genomes of the mouse gut is presented in **Figure 1a**. We combined pre-assembled genomes obtained from public databases and newly-assembled genomes into MRGM. To assemble the novel genomes of the mouse gut microbiome, we collected public WMS data for 838 fecal or cecal samples that have not yet been used for metagenome assembly. Additionally, we generated in-house WMS data from 40 mouse fecal samples. Consequently, we executed a computational pipeline for *de novo* genome assembly for the 878 WMS datasets (**Supplementary Table 1**). We obtained a total of 35,704 new MAGs possessing completeness of ≥ 50%, contamination of ≤ 10%, and quality scores (*Completeness* - 5 * *Contamination*) ≥ 50, and this is equivalent to the medium quality (MQ) according to the minimum information about a metagenome-assembled genome (MIMAG) (46). Next, we combined the newly-assembled genomes with MAGs and isolated genomes from several public databases that included iMGMC (15), MMGC (26), miBC (12), mGMB (13), and PATRIC (27). The quality of MAGs is conventionally evaluated based on their completeness and contamination of redundant and overlapping genomic fragments according to CheckM (25). However, this method cannot detect contamination by non-redundant fragments from unrelated lineages and ultimately results in chimeric genomes. Therefore, we performed an additional quality assessment using the GUNC (28) tool that detects genome chimerism using the homogeneity of contigs and reports the entropy-based metric CSS. We filtered for genomes possessing completeness ≥ 70%, contamination ≤ 5%, and CSS ≤ 0.45 (the default threshold by the software), and this resulted in a total of 58,684 genomes for the MRGM. To count the number of non-redundant genomes, we performed average-linkage hierarchical clustering at a cutoff of 0.001 using the Mash distance (Mash ANI 99.9%). Consequently, we observed that the MRGM contains 46,267 non-redundant genomes for the mouse gut microbiome (genome counts for each step of the pipeline are summarized in **Supplementary Table 2**).

**Figure 1.**
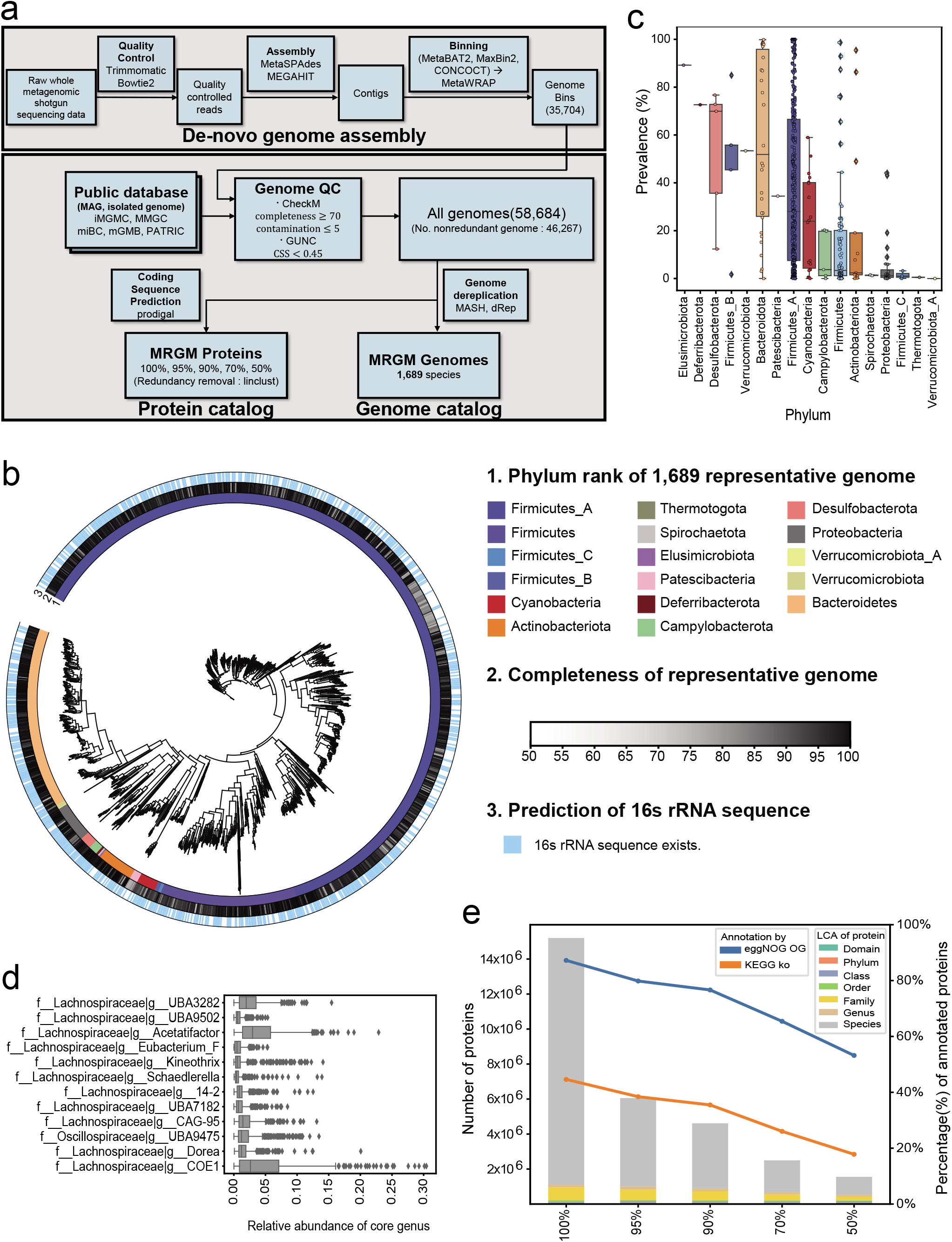
Pipeline and summary information for MRGM. (a) Summary of the bioinformatics pipeline for MRGM (Mouse Reference Gut Microbiome) construction. (b) Phylogenetic tree of 1,689 species of MRGM that summarizes phylum annotation, genome completeness, and availability of 16S rRNA sequence. (c) Prevalence of genera across 878 samples used for MAG (metagenome-assembled genome) assembly is summarized for each phylum. The three major phyla are highlighted by red coloring. (d) Relative abundance of 12 genera that are present in all 878 samples. (e) Summary of MRGM protein catalogs according to clustering with 100%, 95%, 90%, 70%, and 50% sequence similarity. Proportion of functional annotation by eggNOG ortholog group (OG) and KEGG orthology (KO) decreases as the sequence similarity decreases. MRGM also provides information regarding the lowest common ancestor (LCA) for each protein cluster.

We dereplicated 58,684 genomes iteratively to 1,689 species clusters of genomes (**Figure 1b, Supplementary Table 3**). We selected the genomes with the highest quality for each species cluster (species-level genome bins, SGBs) as representatives. Each representative genome of the species was taxonomically annotated using GTDB-Tk (32). We observed only bacterial genomes and no archaeal genome among the 1,689 species, and these were comprised primarily of Firmicutes_A (63.5%), Bacteroides (14.5%), and Firmicutes (11.0%).

Identification of 16S rRNA sequences for each species will allow functional profiling via direct use of the corresponding species genomes and their gene content with amplicon-based metagenome analysis. We thus predicted the 16S rRNA sequence of representative species using the barnnap of Prokka (47). Highly conserved genomic regions such as 16S rRNA genes are typically not efficiently assembled from short-read sequencing data (48). To increase the chances of recovering 16S rRNA sequences, we utilized not only representative genomes but also other conspecific genomes for each species. Nevertheless, we identified 16S rRNA sequences for only 790 (46.8%) out of 1,689 species.

Next, we surveyed the prevalence of genera among the 878 mouse gut microbiomes used for MAG assembly in this study. The three major phyla—Firmicute_A, Bacteroides, and Firmicutes—contain the majority of genera that exhibit widely different prevalence levels across mouse gut microbiomes (**Figure 1c**). We identified 12 genera that were present in all 878 samples (**Figure 1d**). Notably, 11 of 12 ubiquitous genera belong to the *Lachnospiraceae* family that belongs to the Firmicute_A phylum. The *Lachnospiraceae* family members are anaerobic spore-forming bacteria, and they represent the major taxa that are involved in fermenting diverse plant polysaccharides to short-chain fatty acids (SCFAs) (49). Taxonomic members of this family are among the core human gut microbiota that influence human health (50). A previous survey of the genus prevalence based on NCBI taxonomy and 184 samples used for a catalog of mouse gut metagenomes reported the 20 most abundant core genera shared between mice and humans (14). Five of these belonged to the *Lachnospiraceae* family (*Butyrivibrio, Roseburia, Blautia, Coprococcus,* and *Marvinbryantia*). The results of our survey of genus prevalence based on the latest taxonomic classification database (GTDB) and a much-expanded phylogenetic tree for mouse gut microbiota (from 541 species to 1,689 species) suggest that *the Lachnospiraceae* family contains many core taxa for the mouse gut microbiome that may play important roles in generating SCFAs from fibers.

We also cataloged mouse gut microbial proteins by predicting 138,098,166 coding sequences from 58,684 genomes using Prodigal (36). To reduce redundancy for the protein catalog, we clustered protein sequences with a minimum sequence identity of 100%, and this resulted in ~15.2 million non-redundant protein sequences. Many homologous protein sequences exist in the microbiome. Thus, we subsequently clustered protein sequences with similarities of 95%, 90%, 70%, and 50%, and this resulted in ~6.1, ~ 4.6, ~2.5, and ~1.6 million protein sequences, respectively (**Supplementary Table 4**). We performed functional annotation of the proteins using eggNOG-mapper (39). Notably, as the protein similarity decreased, the functional annotation rate decreased (**Figure 1e**). This may be due to the increased proportion of proteins that are specific for mouse gut microbiota and that cannot be annotated by known functions for orthologous groups such as eggNOG (51).

In summary, the MRGM genome catalog provides a total of 58,684 (46,267 non-redundant) genomes possessing completeness ≥ 70%, contamination ≤ 5%, and CSS ≤ 0.45 for 1,689 prokaryotic species. Of these, 790 are also available with 16S rRNA sequences. The MRGM protein catalog provides over 15 million non-redundant protein sequences. This information for the mouse reference gut microbiome is available for free from the MRGM web server (https://www.mbiomenet.org/MRGM/).

### Sequencing depth correlates with MAG quality and genome assembly efficacy for low-abundance microbial taxa in mouse gut

We previously demonstrated the positive effect of sequencing depth on MAG assembly for human gut microbes (10). Microbiota complexity (i.e., the number of species) may affect the efficiency of MAG assembly, and it is much lower in the mouse gut than it is in the human gut. To confirm the relationship between sequencing depth and MAG quality, we generated simulated datasets for different sequencing depths using 10 WMS samples subjected to ultra-deep sequencing (>130 million read pairs). For each of the 10 samples, 0.5, 2.5, 5, 10, 20, 40, 80, and 125 million read pairs were randomly selected. We assembled MAGs for 80 simulated datasets (10 samplesⅹ8 depths) and then compared the quality of conspecific MAGs in two different simulated samples at adjacent sequencing depths. We observed that MAGs from mouse gut metagenomes with greater sequencing depth possessed significantly higher quality in regard to completeness, N50, and contamination (**Figure 2a-c**).

**Figure 2.**
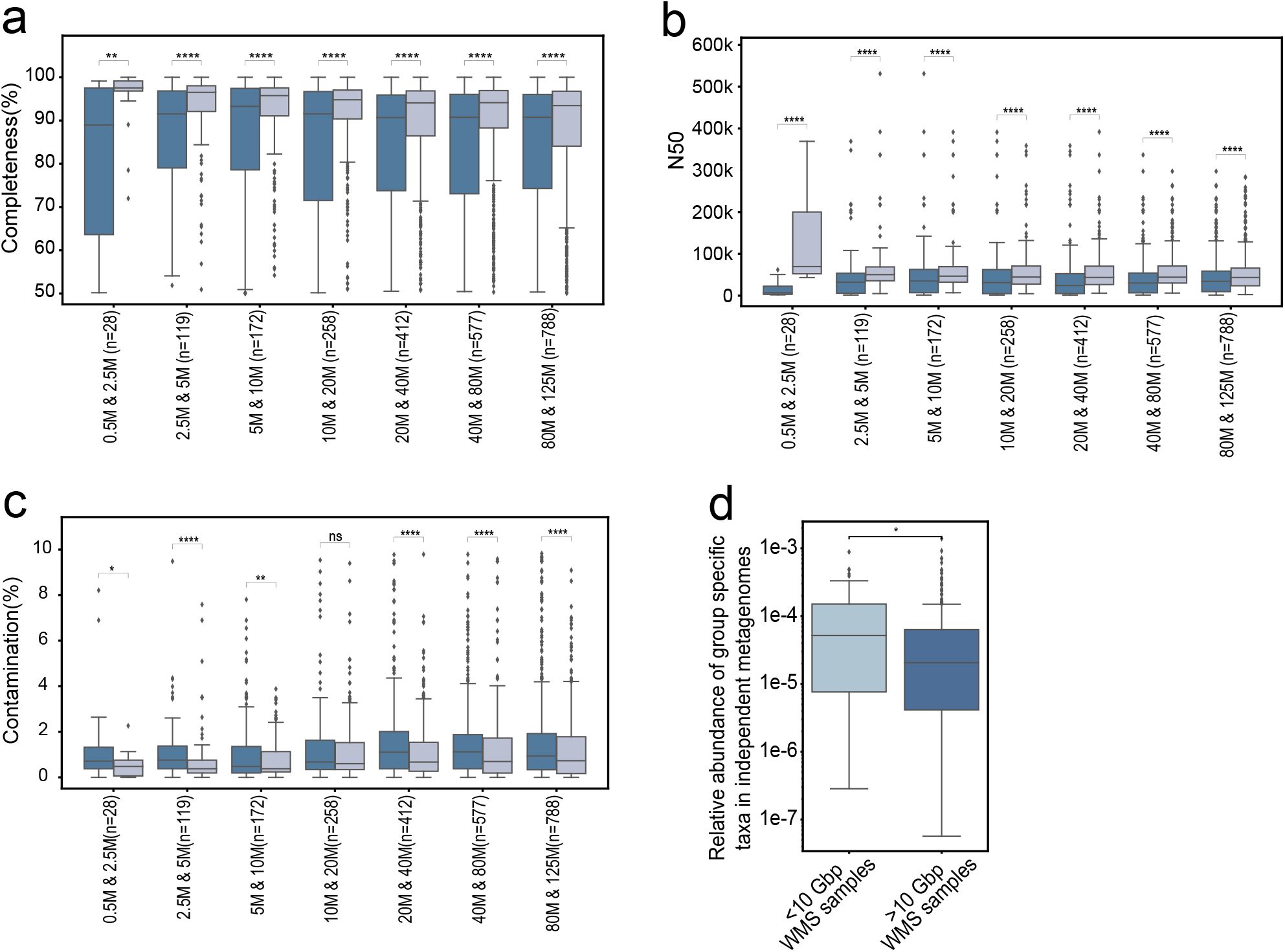
Effect of sequencing depth on MAG assembly. (a-c) Completeness (a), N50 (b), and contamination (c) of assembled genomes according to lower sequencing depth (left box of each column) and greater sequencing depth (right box of each column). (d) The relative abundance of assembled genomes specific for either group of independent fecal samples for different sequencing depths. *P* values were calculated by two-sided Mann–Whitney *U* test (*: *P* <□0.01).

Improved *de novo* genome assembly may enable the reconstruction of MAGs for low-abundance taxa. To test this hypothesis, we divided 378 WMS samples that were used for MRGM construction into two groups based on the sequencing depth range, and these groups included <10 Gbp and >10 Gbp. After filtering out MAGs that also existed in iMGMC or MMGC, we obtained two groups of MAGs that were specific for different ranges of sequencing depth. We then estimated the relative abundance of taxa that were specific for each group in 10 WMS samples possessing the largest sequencing depths (>80 Gbp or >260 million read pairs) of PRJNA603829 (https://www.ebi.ac.uk/ena/browser/view/PRJNA603829) and that were not included in MRGM using Kraken2. We observed a lower range of abundance for taxa that were specifically assembled from WMS samples with higher sequencing depth (>10 Gbp) (**Figure 2d**). These results suggest that deep sequencing may aid in MAG assembly of low-abundance bacterial taxa in the mouse gut.

### CheckM with lineage-specific markers underestimates MAG completeness

A previous catalog, iMGMC, provided mouse gut microbial genomes with MQ. We expected a higher quality of MAGs from the mouse gut microbiome due to its lower complexity compared to that of the human gut microbiome. Thus, we initially used the higher-level criterion for genome quality as defined by MIMAG, where near complete (NC) genomes satisfy the requirements for completeness ≥ 90% and contamination ≤ 5%. However, we found that a genome completeness of ≥ 90% according to CheckM based on lineage-specific markers is too stringent for some taxa. At the order level, the majority of the genomes of the Gastranaerophilales order of Cyanobacteria phylum, the RF39 order of Firmicutes phylum, the TANB77 order of Firmicutes_A phylum, the UBA406B and Peptococcales orders of Firmicutes_B phylum, and the Rs-D84 order of Proteobacteria phylum could not pass the 90% completeness threshold (**Figure 3a**). Furthermore, genomes for the phylum Cyanobacteria and Firmicutes_B exhibited phylum-wide exclusion by completeness ≥ 90% (**Figure 3b**). CheckM estimates the completeness and contamination of MAGs using lineage-specific single-copy marker genes (SCGs). Thus, the CheckM procedure first maps the query MAG into a tree of reference genomes and then selects SCGs specific for the mapped lineage. Thus, misidentification of the lineage of the query MAG may result in an underestimation of completeness. Many new prokaryotic genomes have recently been assembled from metagenome samples and deposited into the public domain. However, the reference genome database for CheckM has not been updated since 2015. Therefore, tree mapping of query MAGs for the recently expanded lineages may result in higher chances of incorrect lineage identification, thus resulting in an underestimation of completeness.

**Figure 3.**
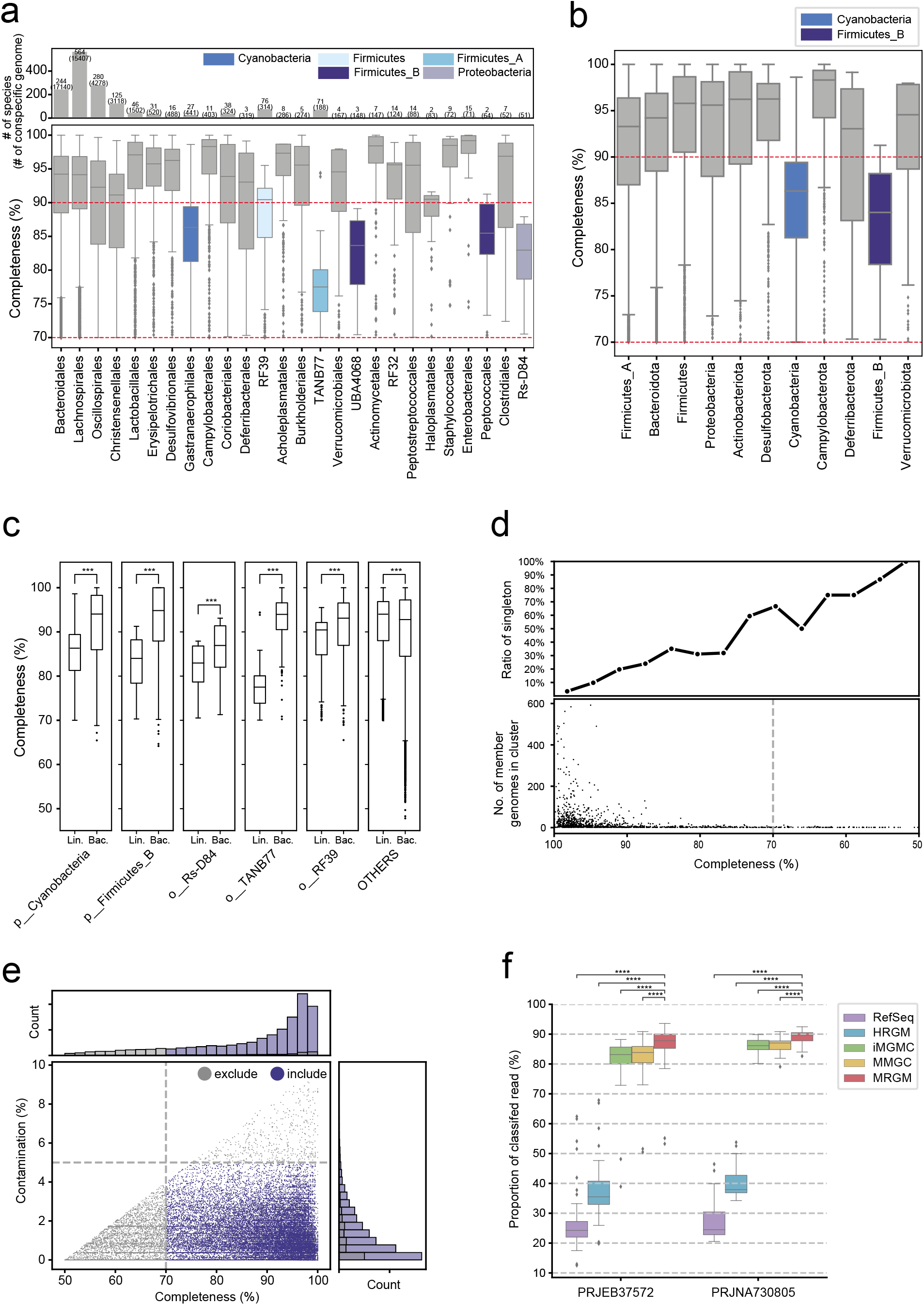
Quality assessment of MRGM genomes. (a-b) Distribution of completeness by CheckM for orders (a) and phyla (b) of MRGM (Mouse Reference Gut Microbiome). MAGs (metagenome-assembled genomes) with completeness ≥ 70%, contamination ≤ 5%, and passing GUNC filtering (CSS < 0.45) were included. (c) Comparison of genome completeness as estimated by lineage-specific markers (Lin.) with those estimated by markers for the entire bacterial species (Bac.). Significance was measured by two-tailed Mann Whitney U test (***, p < 0.001). (d) The ratio of clusters with a single member genome (singleton) for given genome completeness. (e) Summary of estimated completeness and contamination of 58,684 genomes collected from databases and the *de novo* assembly pipeline. Genomes included into the final MRGM are highlighted by blue coloring. (f) Proportion of classified sequence reads from mouse gut WMS (whole metagenome sequencing) data from two published studies by different microbial genome catalogs. Significance was measured by two-tailed Mann Whitney U test (****, p < 0.0001).

To test our hypothesis, we measured completeness using SCGs for all bacterial reference genomes instead of lineage-specific genomes. Notably, genome completeness for the five orders that were predominantly <90% was significantly increased by CheckM evaluation based on bacterial marker genes (**Figure 3c**). For example, majority of genomes for the TANB77 order exhibited completeness between 70% and 80% based on lineage-specific markers, whereas this value was between 90% and 100% based on the bacterial markers. In contrast, genomes for all other orders possessed significantly decreased completeness based on bacterial markers instead of lineage-specific markers. These results suggest that CheckM completeness estimation based on lineage-specific markers works properly for relatively well-established taxa groups but not for uncharacterized ones. We also found that the ratio of species clusters with only a single member genome (singleton) gradually increased as the genome completeness decreased, and genomes with completeness of 50% were comprised predominantly of singletons (**Figure 3d**). Therefore, for the MRGM we used 70% as the completeness cutoff to salvage genomes for the five orders that could be excluded by conventional criterion for the high-quality genome and to avoid too many singletons that are likely to be due to the misassembled MAGs (**Figure 3e**).

### MRGM improves taxonomic classification of sequencing reads

DNA-based taxonomic profiling of metagenomic data relies on the quality and comprehensiveness of the reference genome database (52). The standard database for microbial genomes may lack many of the genomes of prokaryotic species that exist in the mouse gut, and this can lead to false negatives. Genomes that do not exist in the mouse gut environment are also included in the standard databases, and these irrelevant genomes may result in false positives (53). As MRGM is more comprehensive than are previous databases and is specific to the mouse gut microbiome, we expected improved taxonomic classification of metagenome sequencing reads. We used Kraken2 to compare the taxonomic classification of sequencing reads across different prokaryotic genome databases, including RefSeq-based standard database and custom databases constructed from the genomes of iMGMC (15), MRGM, and HRGM (10). We also included a recently released public Kraken2 database from MMGC (26). We assessed the taxonomic classification rate using 86 WMS samples according to single-end sequencing from PRJEB37572 (44) and 30 WMS samples by paired-end sequencing from PRJNA730805 (45) that were not used for the construction of any of the genome catalogs. We observed a significant increase in the classification rate by MRGM compared to that of all other databases (**Figure 3f,** *P* < 0.0001, two-tailed Mann Whitney U test). These results suggest that the expanded microbial genome catalog specific for the mouse gut may improve the efficacy of taxonomic profiling of mouse gut metagenomes.

### Reliable estimation of low-abundance bacterial taxa of the mouse gut requires deep sequencing

A previous study demonstrated a high correlation of overall taxonomic profiles of the human gut microbiome by according to shallow sequencing (0.5-2 million reads) with those obtained by ultra-deep sequencing (2.5 billion reads) (54). Nevertheless, the correlation may not be equally high for low-abundance taxa profiles. Therefore, we sought a minimum sequencing depth to obtain reliable profiles for low-abundance taxa in the mouse gut microbiome. First, we stratified taxonomic features at the domain, phylum, class, order, family, genus, and species level according to mean relative abundance (ranging from 1e-7 to 1 with every ten-fold increase) using 10 WMS samples with the largest sequencing depth (>80 Gbp or >260 million read pairs) of PRJNA603829 (https://www.ebi.ac.uk/ena/browser/view/PRJNA603829) that were not used for MRGM (**Figure 4a**). As the group <1e-7 included only three taxonomic features, we merged it with the group specific for <1e-6. Taxa possessing a relative abundance of <1e-6 accounted for less than 5%, whereas those with <1e-5 comprised 27.44% of the mouse gut microbiome (**Figure 4b**). Therefore, we chose a sequencing depth that could obtain a high correlation of abundance profiles for taxa <1e-5 with those obtained by ultra-deep sequencing. To examine the correlation between taxonomic profiles at different sequencing depths, we generated simulated datasets for 10 different depth ranges (0.5, 1, 2.5, 5, 10, 20, 40, 60, 80, and 125 million read pairs) from each of the 10 WMS samples. Correlations between taxonomic profiles for the 100 simulated datasets were calculated using the Pearson correlation coefficient (*PCC*) (**Figure 4c**). Notably, a shallow sequencing depth, 0.5 million reads, could not achieve a *PCC* > 0.8 for profiles for abundant taxa with a relative abundance of <1e-4, and these accounted for >62% of the mouse gut microbiome (**Figure 4d**). Our analyses suggest that we must sequence >10 million read pairs to achieve a *PCC* > 0.9 for profiles for taxa, including those possessing a relative abundance of <1e-5 that account for >27% of the mouse gut microbiome. We observed similar results with a slightly lower overall correlation based on the Spearman correlation coefficient (*SCC*) (**Figure 4e**).

**Figure 4.**
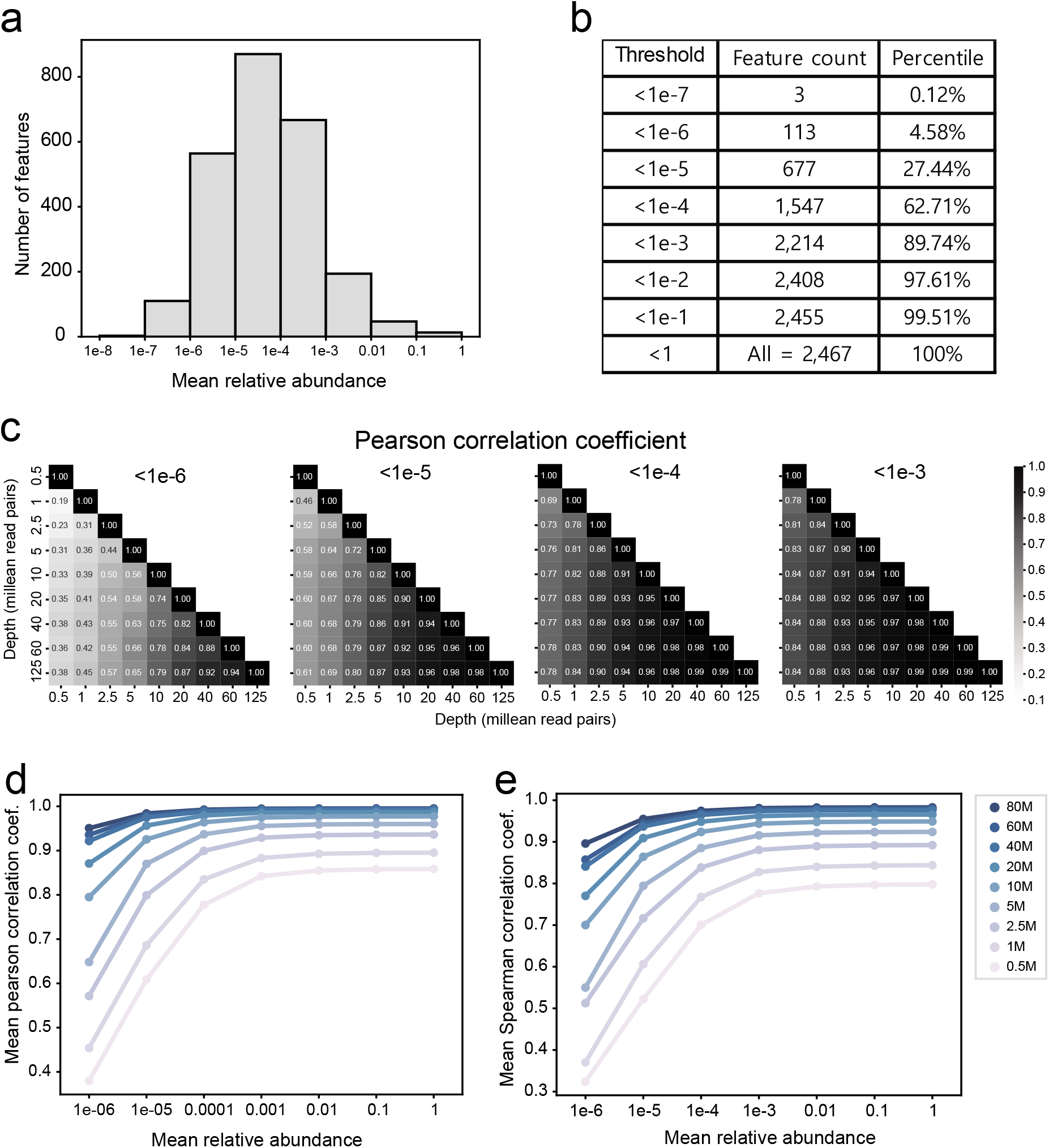
Effect of sequencing depth on the reliability of taxonomic profiles. **(a)**The distribution of taxonomic features over different mean relative abundances. (**b)** The cumulative proportion of taxonomic features at different thresholds of mean relative abundance. (c) Pearson correlation coefficient of the taxonomic profiles between the various sequencing depths (x- and y-axis) for the given mean relative abundance thresholds (title of each heatmap). (d-e) Summary plot of Pearson correlation coefficients (*PCC*) (d) and Spearman correlation coefficients (*SCC*) (d) of the taxonomic profiles at the given sequencing depth and at 80□million reads. The x-axis (the mean relative abundance threshold) indicates the upper boundary of the mean relative abundance.

### Functional capacity of the mouse gut microbiome is largely shared in the human gut microbiome

Notably, although HRGM contains bacterial genomes present in the gut environment, it exhibited an unexpectedly low classification rate for sequencing reads from mouse gut metagenomes (**Figure 3f**). This suggests that bacterial taxa in the gut environment are distinct between humans and mice. A previous study demonstrated that the majority of the KEGG orthology (KO) terms supported by the mouse metagenome catalog are also present in the human metagenome catalog (14), and we could reproduce similar degrees of similarity between MRGM and HRGM in regard to the total KO, Gene Ontology (GO), and carbohydrate-active enzymes databases (CAZy) (42) (**Figure 5a**). Therefore, despite the taxonomic discrepancy in the gut microbiome, the overall functional capacity is largely shared between mice and humans.

**Figure 5.**
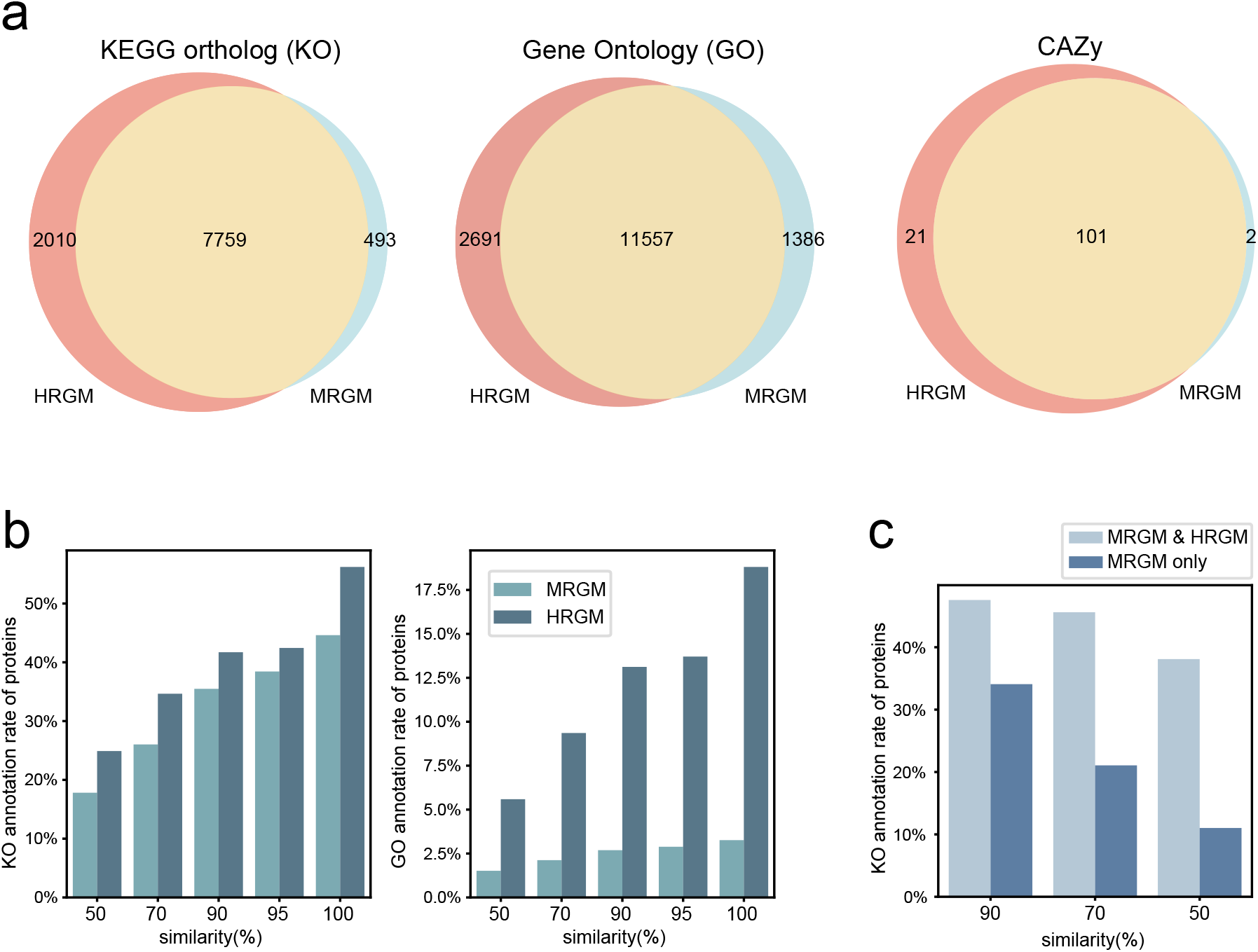
Functional comparison of gut microbiomes between mice and humans. (a) Venn diagram for overlap of total functional terms that annotate gut microbial proteins between MRGM (Mouse Reference Gut Microbiome) and HRGM (Human Reference Gut Microbiome). (b) Comparison of KO and Gene Ontology (GO) annotation rate for protein clusters according to different similarity between MRGM and HRGM. (c) KEGG Orthology (KO) annotation rate for protein clusters shared by both HRGM and MRGM or those specific for MRGM with different similarity.

Next, we compared the functional annotation rates of gut microbial proteins between MRGM and HRGM. We observed that functional annotation rates were generally lower for mouse gut microbial proteins than they were for humans (**Figure 5b**). In particular, the GO annotation rate for mouse gut microbial proteins was less than 3% at all similarity levels. Notably, KO annotation rates for gut microbial proteins specific for mice were substantially lower than were those for proteins that were conserved in humans (**Figure 5c**). These results demonstrate challenges regarding the functional annotation of mouse gut microbial proteins that have no orthologous proteins.

### Only ~10% of mouse gut bacterial species are present in the human gut

We next evaluated the taxonomic overlap between the MRGM and HRGM. To perform a taxonomic comparison of genomes of the same quality, human gut microbial genomes from HRGM were further filtered for CheckM completeness ≥ 70% and GUNC CSS < 0.45. We also excluded archaeal genomes of HRGM, as MRGM contains no archaeal genomes.

Consequently, we used 1,689 species from MRGM and 4,604 species from HRGM for taxonomic comparison. First, we observed different phyla compositions of the gut microbiome in mice and humans (**Figure 6a**). For example, the Firmicutes_A phylum possessed 1,073 of 1,689 (63.5%) MRGM species; however, only 1,884 (40.9 %) were present in the same phylum as HRGM. Actinobacteriota is the second largest phylum of the human gut microbiome, whereas only 49 species of MRGM (2.9 %) belong to the same phylum as mouse gut bacteria.

**Figure 6.**
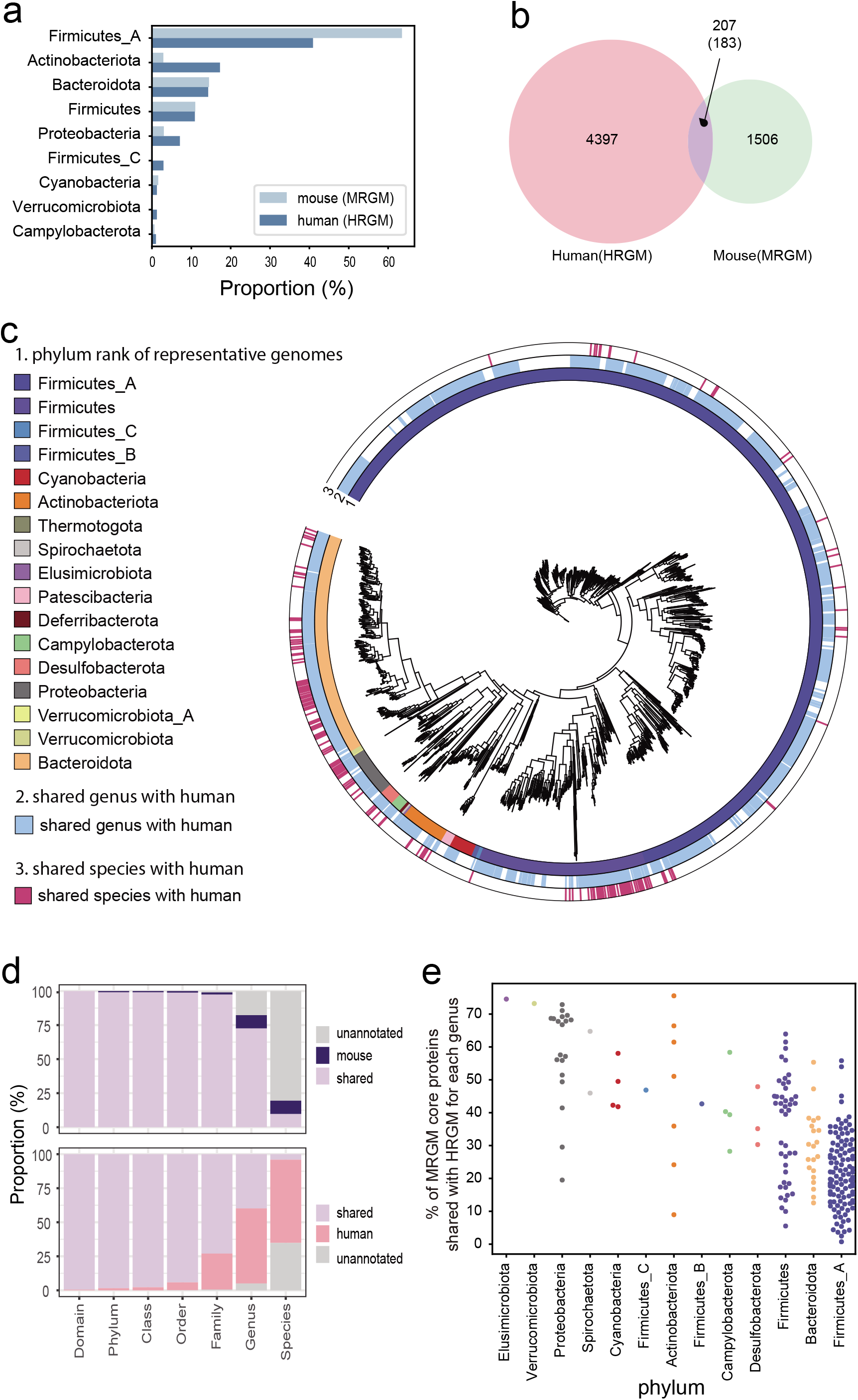
Taxonomic comparison of gut microbiomes between mice and humans. (a) Proportion of each phylum in the mouse and human gut microbiomes. (b) Only 183 of 1,841 representative species (~10%) of gut bacterial species are shared between humans and mice. (c) A total of 1,841 representative genomes were annotated by phylum annotation (1^st^ layer from the inside). If the genus and species were shared with HRGM (Human Reference Gut Microbiome), it was annotated by blue and pink (2^nd^ and 3^rd^ layer). By phylum, the shared pattern was different. In Bacteroidota, numerous species were shared, whereas in Firmicutes_A, only a small number of species were shared. (d) Proportion of shared or specific taxonomic groups between mice (top) and humans (bottom) for each of the taxonomic classification levels. (e) Percentage of core genes for the same genus between HRGM and MRGM.

Taxonomic overlap may be simply measured based upon the total number of shared taxonomic groups for each classification level between the two gut microbiomes. However, we observed that 1,360 of 1,689 (80%) of MRGM species representative genomes possessed no assigned species groups, and the majority of them were taxonomically annotated at the genus level and above. Thus, simple counting of shared taxonomic groups is not feasible for species-level taxonomic comparisons. Therefore, we used ANI for taxonomic comparisons at the species level. We counted a pair of gut bacterial species genomes from humans and mice as shared species if ANI ≥ 95. We observed that only 183 of 1,689 (10.8%) MRGM species and 207 of 4,604 (4.5%) HRGM species were shared between humans and mice (**Figure 6b**) (**Supplementary Table 5**). The majority of the shared bacterial species belonged to the phyla Proteobacteria, Firmicutes, and Bacteroidota (**Figure 6c**).

### Gut microbes for same genus typically share a minor portion of core gene content between mouse and human

Conversely, we observed high overall taxonomic similarity at the genus level and a higher level of taxonomic classes in which most of the MRGM genomes possessed HRGM genomes for the same taxonomic groups (**Figure 6d**). For example, 220 of 278 (79.1%) MRGM genera possessed genomes for the same genus in HRGM (**Supplementary Table 6**). Particularly, MRGM Firmicute_A phylum that does not possess a large number of species overlaps with HRGM exhibited numerous shared genera (**Figure 6c**). Nevertheless, bacteria for the same taxonomic groups in the same body site could be dissimilar in their core gene content, and in turn, they could carry different functional repertoires. Therefore, we first identified ‘genus core’ genes for protein groups according to 50% similarity that are conserved among ≥90% of non-redundant genomes for each genus, and we then measured proportion of MRGR core genes shared with HRGM for each genus (**Materials and Methods**). Notably, the majority of the genera for the three major phyla, Firmicute_A, Bacteroidota, and Firmicutes of MRGM possessed <40% of core genes that overlapped with their HRGM counterparts (**Figure 6e**). Among all 220 shared genera of MRGM, 155 (70.5%) and 119 (54.1%) genera possessed <40% and <30% of core genes, respectively (**Supplementary Table 7**). These results indicate that many intestinal commensal bacteria of the same genus may play distinct roles in the mouse and human gut. The results also suggest that mouse gut microbiome research may not be easily translated into human microbiome research by genus-level taxonomic commonality.

## DISCUSSION

In the present study, we constructed a reference gut microbiome for mouse, MRGM, which provides an expanded catalog of genomes and proteins by MAG assembly from 878 new samples with deep metagenomic sequencing. We demonstrated a significantly improved classification rate of metagenomic sequencing reads by MRGM compared to those of the standard database of the prokaryotic genome and of previously published databases of mouse gut microbial genomes. These results indicate that gut microbial dark matter was effectively uncovered through the additional MAG assembly in the present study.

Our work is distinct from earlier studies on the construction of the mouse gut microbiome catalog in several aspects. First, we found that the current database of lineage-specific markers underestimated the completeness of MAGs for many of the relatively uncharacterized taxa. Thus, we cataloged MAGs using a threshold of 70% for CheckM completeness. Genomes for unculturable bacteria are relatively uncharacterized, and thus, the majority of the genomes with completeness between 70% and 90% may represent unculturable bacterial taxa. Given that the majority of intestinal commensal species are uncultivable, this new criterion for MAG completeness will provide more opportunities for exploring gut microbial dark matter. Second, we found that reliable profiles for low-abundance taxa require deep sequencing rather than shallow sequencing. Therefore, we recommend sequencing >10 million read pairs for taxonomic profiling based on metagenomic sequencing. Finally, we demonstrated a high overlap of genera; however, there was still a large functional discrepancy between mouse and human gut microbiota. Findings from mouse gut microbiota are often transferred to human gut microbiome studies, and this often occurs via shared genera. The results of this study suggest that functional translations of metagenomics research in the mouse gut into the human gut microbiome by genus-level taxonomic commonality require caution.

## DATA AVAILABILITY

Raw metagenomic sequencing data are available from the NCBI Sequence Read Archive (SRP335854). MAGs assembled from 878 mouse gut metagenomes were also deposited on the web server. By accessing the web server (https://www.mbiomenet.org/MRGM/), users can browse and download all genomes for 1,689 representative species and also their annotations and metadata, including taxonomy, genomic content, and genome statistics. The five classes of protein catalogs and 16S rRNA sequences are also provided with their functional annotation and taxonomic origin.

